# Quiescent OXPHOS-high triple-negative breast cancer cells that persist after chemotherapy depend on BCL-XL for survival

**DOI:** 10.1101/2025.08.21.671546

**Authors:** Slawomir Andrzejewski, Marie Winter, Leandro Encarnacao Garcia, Olusiji Akinrinmade, Francisco D.M. Marques, Emmanouil Zacharioudakis, Anna Skwarska, Julio Aguirre-Ghiso, Marina Konopleva, Guangrong Zheng, Susan Fineberg, Daohong Zhou, Evripidis Gavathiotis, Tao Wang, Eugen Dhimolea

**Affiliations:** Department of Molecular Pharmacology, Albert Einstein College of Medicine, Bronx, NY, USA; Montefiore Einstein Comprehensive Cancer Center, Albert Einstein College of Medicine, Bronx, NY, USA; Cancer Dormancy Institute, Albert Einstein College of Medicine, Bronx, NY, USA; Department of Biochemistry, Albert Einstein College of Medicine, Bronx, NY, 10461, USA; Department of Medicine, Albert Einstein College of Medicine, Bronx, NY, USA; Department of Cell Biology, Albert Einstein College of Medicine, Bronx, NY, USA; Department of Oncology, Albert Einstein College of Medicine, Bronx, NY, USA; Ruth L. and David S. Gottesman Institute for Stem Cell Research and Regenerative Medicine, Albert Einstein College of Medicine, Bronx, NY, USA; Institute for Aging Research, Albert Einstein College of Medicine, Bronx, NY, USA; Marilyn and Stanley M. Katz Institute for Immunotherapy for Cancer and Inflammatory Disorders, Albert Einstein College of Medicine, Bronx, NY, USA; Department of Medicinal Chemistry, College of Pharmacy, University of Florida, Gainesville, FL, USA; Department of Pathology, Montefiore Medical Center, New York, NY, USA; Department of Biochemistry and Structural Biology, University of Texas Health San Antonio, San Antonio, TX; Department of Epidemiology & Population Health, Albert Einstein College of Medicine, Bronx, NY, USA

**Keywords:** Quiescence, Oxidative phosphorylation, BCL-XL, Chemotherapy, Triple-negative breast cancer

## Abstract

The persistent residual tumor cells that survive after chemotherapy are a major cause of treatment failure, but their survival mechanisms remain largely elusive. These cancer cells are typically characterized by a quiescent state with suppressed activity of MYC and MTOR. We observed that the MYC-suppressed persistent triple-negative breast cancer (TNBC) cells are metabolically flexible and can upregulate mitochondrial oxidative phosphorylation (OXPHOS) genes and respiratory function (“OXPHOS-high” cell state) in response to DNA-damaging anthracyclines such as doxorubicin, but not to taxanes. The elevated biomass and respiratory function of mitochondria in OXPHOS-high persistent cancer cells were associated with mitochondrial elongation and remodeling suggestive of increased mitochondrial fusion. A genome-wide CRISPR editing screen in doxorubicin-persistent OXPHOS-high TNBC cells revealed BCL-XL gene as the top survival dependency in these quiescent tumor cells, but not in their untreated proliferating counterparts. Quiescent OXPHOS-high TNBC cells were highly sensitive to BCL-XL inhibitors, but not to inhibitors of BCL2 and MCL1. Interestingly, inhibition of BCL-XL in doxorubicin-persistent OXPHOS-high TNBC cells rapidly abrogated mitochondrial elongation and respiratory function, followed by caspase 3/7 activation and cell death. The platelet-sparing proteolysis targeted chimera (PROTAC) BCL-XL degrader DT2216 enhanced the efficacy of doxorubicin against TNBC xenografts *in vivo* without induction of thrombocytopenia that is often observed with the first-generation BCL-XL inhibitors, supporting the development of this combinatorial treatment strategy for eliminating dormant tumor cells that persist after treatment with anthracycline-based chemotherapy.

## INTRODUCTION

Cytotoxic chemotherapy remains a cornerstone in the treatment of many highly lethal cancers, including triple negative breast cancer (TNBC). However, a fraction of tumor cells can survive chemotherapy as viable “seed” for the eventual relapse. These residual cancer cells are a crucial link in the disease evolution as they represent the reservoir of malignant cells from which future genetic, epigenetic and other forms of treatment resistance emerge. The eradication of persistent cells could achieve prolonged remissions or cures. Recent work by us [1,2] and others [3,4] in multiple cancer types has shown that chemotherapy-persistent cancer cells in preclinical models and in patients can respond to drug-induced cytotoxic stress by adopting a reversible biosynthetically/proliferatively quiescent state, distinct from baseline tumors and not genetically selected [1,2,4]. This rapid transition of residual cancer cells to chemo-tolerant quiescence is enabled by suppression of MYC [1,2] and MTOR [1,3], which are global drivers of cell growth and biosynthesis in proliferating cells [5]. In particular, inactivation of MYC induces a biosynthetically paused state with reduced redox stress and attenuated apoptotic priming, which partially mitigates drug-induced cytotoxicity [1-3]. Interestingly, this drug persistence mechanism mimics the “survival-mode” transcriptional reprogramming in embryonic diapause [6], a reversible quiescence stage in early embryos driven by adaptive MYC suppression in response to environmental perturbagens and physiological factors [7].

Anthracyclines (e.g. doxorubicin) and taxanes (e.g. docetaxel) are commonly used for the treatment of TNBC in the neoadjuvant and adjuvant settings [8]. Anthracyclines are nucleic acid intercalators that induce cancer cell death by causing DNA damage which eventually leads to activation of apoptosis [9]. When the anthracycline-induced DNA damage is not sufficient for reaching the apoptotic threshold, the residual cancer cells can enter a G0/G1 quiescent state and activate DNA damage repair mechanisms [1,10]. Distinctly, taxanes are mitotic spindle poisons that are thought to kill cancer cells during mitosis through induction of multipolar spindles and chromosomal instability [11]. Taxane-persistent cancer cells can evade drug-induced mitotic cell death by exiting mitosis (a phenotype referred to as “mitotic slippage” [12]) and entering a diapause-like quiescence until proliferation resumes [1,13]. Both anthracycline- and taxane-persistent cancer cells presumably resume proliferation after drug-induced cellular damage has been repaired.

While the transient dormant state after treatment contributes to drug tolerance, recent published work has demonstrated that persistent cells can also evade cell death by rewiring their metabolism [14,15]. For instance, detoxifying enzymes such as ALDH and GPX4 can be essential for the survival of drug-persistent cancer cells and render residual tumors sensitive to their inhibition [16-19]. More notably, increased OXPHOS function has been linked to reduced drug sensitivity in breast and other cancers [20-25]. For instance, cytarabine-persistent residual acute myeloid leukemia (AML) cells increase their mitochondrial mass, consistent with a high OXPHOS state [26]. Upregulation of peroxisomal fatty acid β-oxidation (FAO) and a metabolic shift from glycolysis to oxidative respiration has also been recently reported in drug-tolerant persistent cells in lung cancer, melanoma and AML [4,27,28]. An increase in mitochondrial respiratory function in response to treatment has been reported in several other solid and hematological cancers [20,22-25]. The increasing recognition of the role of OXPHOS in cancer has prompted the development of OXPHOS inhibitors with excellent selectivity against electron transport chain (ETC) complex I [21,29]. Preclinical evaluation of these OXPHOS inhibitors had shown promising results in several cancer models, including chemotherapy-persistent OXPHOS-high TNBC [20]. Despite their target selectivity and significant therapeutic potential, the advancement of OXPHOS inhibitors in the clinic has been hampered by high levels of adverse effects likely due to the inhibition of cellular respiration in normal cells [30,31], highlighting the need for alternative approaches targeting the quiescent OXPHOS-high persistent cancer cell state.

Here we examined the mitochondrial OXPHOS-high state in MYC-suppressed chemotherapy-persistent TNBC cells in preclinical models and patients and used genetic and pharmacological tools to interrogate its therapeutic vulnerabilities. We observed that the quiescent persistent cancer cell state is metabolically heterogenous across chemotherapeutic classes and cellular models. Notably, the activation of OXPHOS in quiescent post-treatment cancer cells is associated with acquired dependency on BCL-XL and can be targeted therapeutically using a proteolysis targeted chimera (PROTAC) based approach with minimal toxicity.

## MATERIALS AND METHODS

### Cell culture

TNBC cell lines MDAMB-231, SUM159, HCC1806 were obtained from ATCC and maintained in DMEM (Dulbeco’s Minimal Essential Medium, Gibco) supplemented with 10% FBS (Fetal Bovine Serum), 40 UI/mL penicillin (Gibco), 40 μg/mL streptomycin (Gibco) at 37°C in 5% CO_2_ humidity-controlled atmosphere. MDA-MB-468 TNBC cell line (from ATCC) was cultured in RPMI 1640 culture medium supplemented with 10% FBS, and 40 UI/mL penicillin, 40 μg/mL streptomycin at 37°C in 5% CO_2_ humidity-controlled atmosphere. All cell lines were stably transfected with lentiviral vector expressing luciferase and mCherry (FUW-Luc-mCherry-puro, Addgene #183685).

### Generation of drug-persistent cells

The cancer cells were plated in 6-well low-adherence plates (Corning, 1 million cells/cm^2^) and treated with DMSO, or 150 nM or 250 nM doxorubicin (Selleckem) or 15 nM docetaxel (Selleckem) for 2 hours at 37°C. After the pulse treatment, the cells were washed with PBS, followed by centrifugation (5min, 300*xg*) and plating in 10-cm dish (100,000 cells/cm^2^) for 48h or 72h. The quiescent state of the surviving cells was confirmed by cell cycle analysis using flow cytometry. The resulting quiescent drug-persistent cells were used for subsequent experiments.

### In vitro drug response assays

Chemotherapy-persistent and chemo-naïve cells were seeded (1,500 cells/well) in 384-well plates (Corning) 24 hours before the treatment using cell culture medium containing 5% luciferin. The evaluated drugs were added the next day and cell viability is evaluated every 24 hours during 72-96 hours treatment using a plate reader (SynergyH1, Biotek) and bioluminescence as readout.

### Western blot

Cells were washed with cold PBS and lysed in Radio Immuno Precipitation Assay (RIPA) buffer (New England Biolabs) containing protease and phosphatase inhibitor (Thermo Scientific, 78446). Cell lysates were centrifuged and supernatants were collected. Protein concentration was measured using a BCA Protein Assay Kit (Thermo Fisher Scientific). Protein samples were mixed with Laemmli buffer(Bio-Rad) and loaded (5 μg/well) on 4% to 12% SDS-PAGE gels, followed by transfer to PVDF membranes. Membranes were blocked in 5% milk and incubated with primary antibodies at 4°C overnight. After washing with PBST, membranes were incubated with the appropriate secondary antibodies for 2 hours at room temperature. Membranes were incubated with SuperSignal West Femto Stable Peroxide & Luminol/Enhancer (Thermo Fisher Scientific) and developed using the ECL Detection System (Thermo Fisher Scientific). The following antibodies were used: BCL-XL (66020-1), MFN2 (82673-2-RR), anti-rabbit antibody (A8914), anti-mouse antibody (A10516).

### RNA sequencing

Chemotherapy-persistent or chemo-naïve cell pellets were harvested during the proliferation pause phase (3-5 days after the suspension treatment). RNA was extracted using the RNAeasy kit (Qiagen, 74004) following the manufacturer’s instructions. Briefly, the cells were initially counted and then lysed with the RLT buffer supplemented with 5% β-mercaptoethanol. ERCC RNA Spike-In Mix (1/10 dilution, Thermo Fisher Scientific, 4456740) was added in each condition to adjust for the cell number. The extracted RNA was measured using Nanodrop^TM^ spectrophotometers (Thermo Fisher Scientific, ND2000). RNA integrity was assessed using Qubit fluorometer (Thermo Fisher). Libraries were prepared using a library prep kit (Illumina) and sequenced using Illumina NextSeq 500 Next Gen with 150bp paired-end reads (Novogene). Trimmed raw reads were aligned to the reference genome GRCh38 using STAR [56,57]. Reads are counted for each gene with featureCounts (reference), and differentially expressed analysis was performed in DESeq2 [58]. In addition to the results, Gene set enrichment analysis (GSEA) and ingenuity pathway analysis (IPA) were also performed.

### Genome-wide CRISPR-editing screen

MDA-MB-231 cells that stably expressed the Cas9 DNA endonuclease were transfected with the Brunello lentiviral LentiCRISPR v2 library (Addgene #73178-LV) at a MOI of 70% in medium containing polybrene (8µg/mL) for 12h, similarly to our previous studies [59]. The transfected cells were divided in 2 groups: the baseline group and the test group. The cells in the baseline group were store in dry pellet at -80°C 1 week after the transfection. The cells in the test group were treated for 2h in suspension with doxorubicin (125 nM) or docetaxel (10 nM) or nothing (naïve cells). Twenty-four hours after lentivirus washout, the cells were selected through treatment with puromycin (1µg/mL) for 2 days. Baseline samples were collected and stored as cell pellets immediately after puromycin selection was completed. The remaining transfected cells were treated with DMSO or 250nM doxorubicin (2h pulse treatment as described above). The cells were collected after 2 weeks the gDNA was extracted using the QIAamp Blood Maxi kit according manufacturer’s instructions. The sgRNA loci were selectively amplified by PCR using the NEBNext Ultra II Q5 Master Mix (NEB, #M0544), P5 primers mix (500 nM), P7 primer (500 nM). Each replicate was barcoded with a specific P7 primer which also contained the Illumina adaptor sequences. PCR reactions were pooled and purified using AMPure XP magnetics beads (Beckman coulter #A63880). The quantity of gDNA obtained after PCR purification was measured by Qubit. Multiplexed samples were then sequenced using an Illumina NextSeq 500 (Novogene), allowing ∼4x108 individual reads per multiplexed sample.

Relative sgRNA abundance was performed by applying one-sided test for enrichment and depletion of the sgRNAs and sgRNA rank aggregation for each gene using the MaGECK-RRA algorithm and default parameter settings [60]. Analysis was performed for three readcount normalization techniques (total, median and non-targeting readcount normalization). The non-targeting sgRNAs were used as control distribution for the rank aggregation procedure based on the RRA algorithm.

### BH3 profiling

BH3 profiling was performed as previously described [61]. Briefly, alamethicin (25 µM), CCCP (10 µM), and the synthetic peptides BID (final concentrations of 10–0.1 µM) or HRK-y (final concentrations of 10, 1, and 0.5 µM) were added to the JC1-MEB staining solution (150 mM mannitol, 10 mM HEPES-KOH, 50 mM KCl, 0.02 mM EGTA, 0.02 mM EDTA, 0.1% BSA, 5 mM succinate, pH 7.5) in a black 384-well plate. Approximately 40,000 Doxo-P or chemo-naïve MDAMB-231 cells in single-cell format were suspended in 15 µL JC1-MEB staining solution. Cells were incubated for 10 minutes at room temperature in the dark before being dispensed in the 384-well plates containing the peptides. JC1 fluorescence was then monitored using a TECAN M1000 microplate reader (excitation: 545 nm; emission: 590 nm) at 30°C, every 15 minutes for a total of 3 hours. Mitochondrial depolarization was quantified by calculating the area under the curve using GraphPad Prism software (version 8.0.1.244; GraphPad Software, La Jolla, CA, USA).

### Mitochondrial network analysis

Mitochondrial structures were stained using the mitochondria-specific fluorescent dye MitoTracker CMXRos (Thermo Fisher Scientific, M7514). Chemotherapy-persistent and chemo-naïve cells were seeded in clear-bottom 96-well plates (Corning) at approximately 60% confluency the day before. Following treatment, cells were washed twice and incubated in serum-free, pre-warmed medium containing MitoTracker CMXRos at 2 µM for 40 minutes at 37°C in a humidity-controlled atmosphere with 5% CO_2_ and protected from light. After incubation, cells were washed twice and maintained in serum-free, pre-warmed medium at 37°C. Image acquisition was performed using an ECHO microscope (Evolve) with a 10x objective, capturing 10 images per condition. Raw images were processed using ImageJ with the MiNA plugin [62] for mitochondrial network analysis and ridge detection [63], following the authors’ instructions. At least 60 cells were analyzed for each experimental condition. **Caspase activation staining**

The cells were stained using CellEvent™ Caspase-3/7 Detection Reagent (FITC fluorescence; Thermo Fisher Scientific, #C10430), following the manufacturer’s instructions. The day before the assay, persistent and naïve cells expressing mCherry were seeded in 96-well plates at approximately 50% confluency in culture medium. On the day of the experiment, cells were incubated with the CellEvent™ Caspase-3/7 Detection Reagent (10X working solution) and simultaneously treated with either vehicle (untreated), A-1155463, or the mitochondrial complex I inhibitor IACS-010759 (Sigma) at a final concentration of 10 µM. Cells were incubated for at least 1 hour at 37°C in a humidified atmosphere with 5% CO_2_ and protected from light. Caspase-3/7 activation was evaluated at 1, 3, 6, and 24 hours post-staining using fluorescence microscopy (ECHO, Evolve). For each condition, 10 images were acquired per well, across 3 independent wells, using a 10x objective with Texas Red and FITC channels. Raw images were processed using ImageJ. The total number of cells was quantified using a cell-counting plugin in the Texas Red channel, and cells positive for Caspase-3/7 activation were manually counted on the FITC channel. For each time point, the percentage of Caspase-3/7-positive cells was calculated as the ratio of FITC-positive cells to the total number of cells.

### Oxygen consumption rate (OCR) measurement

Oxygen Consumption Rate (OCR) was measured using the XFe96 Seahorse analyzer (Seahorse Bioscience, Billerica, MA, USA) as previously reported [64]. Briefly, cells were suspended in DMEM (Merck, Darmstadt, Germany) with L-glutamine (2 mM), glucose (10 mM) and pyruvate (1 mM) for oxygen consumption and seeded at 25 × 10^4^ cells·100 µL^-1^·well^-1^. Each port was loaded with 20 µL (XFe96) inhibitor diluted in OXPHOS medium. The OCR was assessed at baseline and after injection of each of the following molecules: oligomycin A (1µM), FCCP (0.2 and 0.3 µM), rotenone and antimycin A (1 µM each). To evaluate the impact of BCL-XL on OCR, after the washes with Seahorse medium, the cells were treated with 10 μM of Bcl-XL inhibitor A-1155463 (Selleckem) 1 hour before the Seahorse experiment and maintained under Bcl-XL inhibition during the experiment. After the Seahorse assay the cells were stained using Hoestch33342 (1 mM; Invitrogen) for 30 minutes in the dark at room temp. The staining intensity was measured using the SynergyH1 (Biotek) plate reader and the readout values were used as surrogate for the cell number to normalize the Seahorse output data.

### Animal studies

The in vivo experiments were conducted in accordance with the experimental and ethical guidelines of the Institute of Animal Studies of the Albert Einstein College of Medicine. MDAMB-231 cells were suspended in medium containing 50% matrigel and injected subcutaneously in both flanks of NU(NCr)-Foxn1nu mice (10^6^ cells per injection). When the average tumor diameter reached approximately 0.5 cm, mice were separated into treatment vs. control groups as indicated and treated biweekly (intraperitoneal injections) with either DMSO, doxorubicin (3mg/kg), DT2216 (15mg/kg, provided by Dialectic Therapeutics), or doxorubicin plus DT2216. Tumor growth and response to treatment was quantified by longitudinal tumor size measurements and *in vivo* bioluminescence measurements. Statistical analyses (GraphPad Prism 8.2.0) compared all tumor size values between treatment vs. vehicle cohorts via two-way analysis of variance (ANOVA) with correction for multiple comparisons (Sidak test, default option of the statistical package). Platelets were measured after the last drug injection using a standard automated hematology analyzer.

### Statistical analysis

Data are presented as the mean values ±standard deviation, from at least 3 independent experiments. Statistical significance was measured with GraphPad Prism software (GraphPad software ver 8.0.1.244, Inc., La Jolla, CA, USA) using the unpaired two-tailed student’s *t*-test, or ANOVA test with a *p* value of 0.05 or less considered as statically significant.

## RESULTS

### Mitochondrial respiration is upregulated in doxorubicin-persistent TNBC

To determine the levels of mitochondrial OXPHOS in cancer cells that persist after chemotherapy, we initially examined the transcriptional profiles of quiescent, MYC-suppressed, tumor cells from breast cancer patient samples collected after neoadjuvant chemotherapy (NAC) of combined anthracycline and taxane treatment [32]. As expected, these residual tumor cells had suppressed transcriptional modules related to proliferation and biosynthesis. Interestingly, despite the global transcriptional suppression in this quiescent cell state, a large proportion of the MYC-suppressed residual tumors had increased expression of mitochondrial OXPHOS genes (“MYC-low/OXPHOS-high” cell state, **Figure 1A**), largely reflecting the components of the ETC complexes. The patients with OXPHOS-high post-NAC residual tumors were not restricted to a single breast cancer subtype, but in general tended to have baseline (pre-treatment) tumors with more aggressive clinical features, with many being TNBC. Because increased OXPHOS has been linked with reduced drug sensitivity in breast and other cancers [20-25], we reasoned that targeting this cancer cell state could enhance the efficacy of treatment. To dissect this tumor cell state, we first examined whether the OXPHOS-high transcriptional signature can be modeled in chemotherapy-persistent TNBC cell lines grown *in vitro*. Pulse treatment (2h) of MDAMB-231 cells with clinically relevant concentrations of doxorubicin or docetaxel induced fractional killing, followed by proliferative arrest of the surviving chemotherapy-persistent cells ∼48h after drug exposure (**Figure S1A**). We compared the transcriptional profiles of docetaxel-persistent (**Dtx-P**) and doxorubicin-persistent (**Doxo-P**) cells with their untreated (**chemo-naïve**) counterparts. The Dtx-P cells had suppressed MYC activity and downregulated OXPHOS genes (**Figure 1B**), similar to the “MYC-low/OXPHOS-low” residual tumors in the clinical dataset (**Figure 1A)**. Interestingly, Doxo-P cancer cells also had suppressed Myc activity and overall proliferation modules, but upregulated OXPHOS transcriptional signature (**Figure 1B**), similar to the respective subset of “MYC-low/OXPHOS-high” post-NAC patient tumors (**Figure 1A)**. To determine the effect of OXPHOS gene upregulation on mitochondrial biogenesis and function, we microscopically quantified the mitochondrial length and area (“footprint”) in the quiescent chemotherapy-persistent MDAMB-231 cells (**Figure 1C)**. Qualitative and quantitative microscopic analyses determined that the quiescent OXPHOS-high Doxo-P (but not the OXPHOS-low Dtx-P) cells have increased mitochondrial length, overall footprint and network branching (**Figure 1D)**, which are morphological changes associated with increased mitochondrial fusion and shown to increase OXPHOS function and attenuate apoptosis [22,33,34].

**Figure 1.**
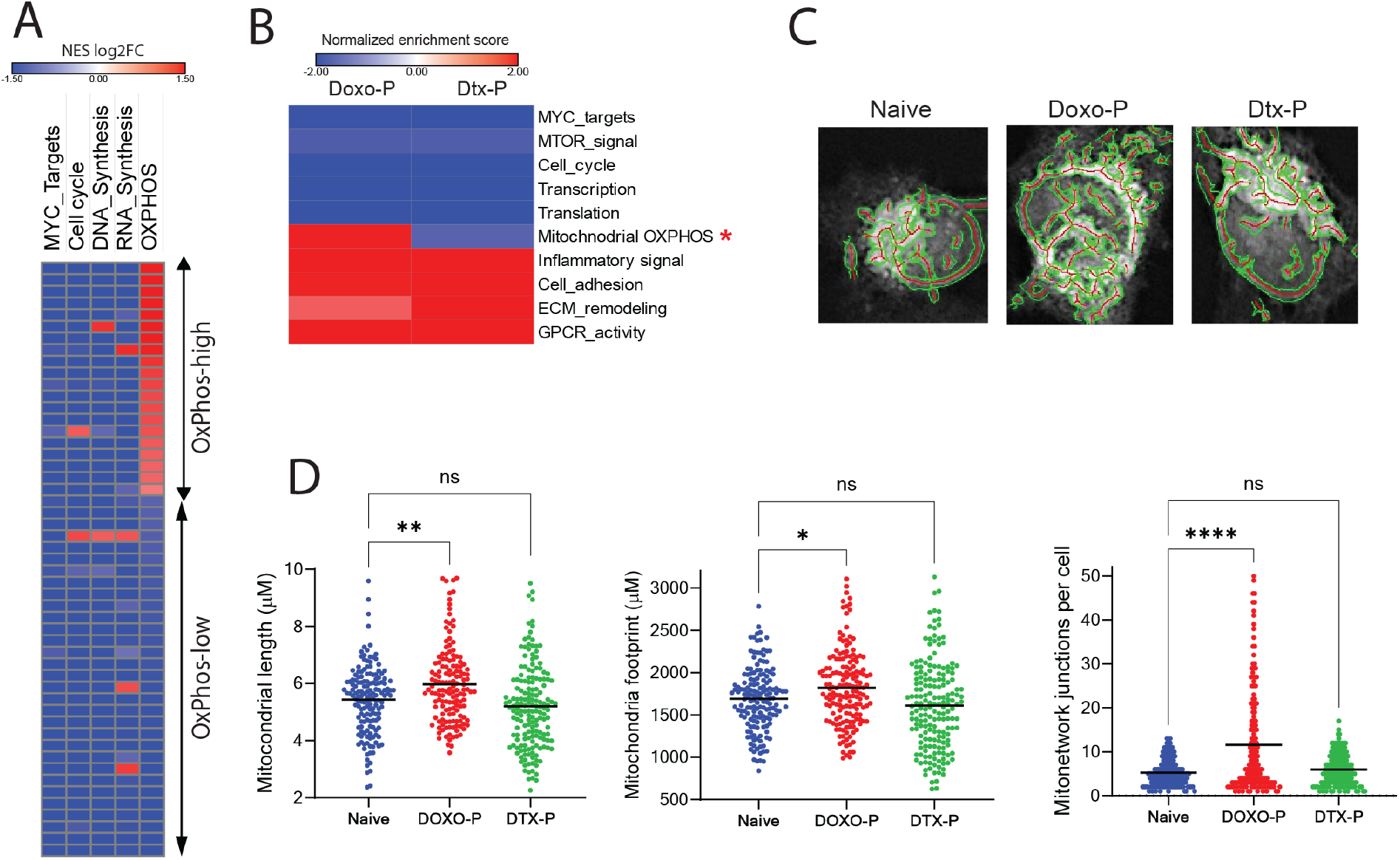
Altered mitochondrial biogenesis and structure in chemotherapy-persistent TNBC cells. **A)** Bulk RNA-seq gene set enrichment analysis (GSEA) of quiescent residual vs. baseline breast tumor cells after chemotherapy in PROMIX trial patient dataset (GSE87455; each row represents a patient). **B)** GSEA analysis of transcriptional changes in MDAMB-231 cells surviving treatment with doxorubicin (Doxo-P) or docetaxel (DTX-P), compared to chemo-naïve counterparts. OXPHOS transcriptional module is marked with asterisk **C)** Mitochondrial network in individual chemo-naïve, Doxo-P and Dtx-P MDAMB-231 cells visualized using the Mito Tracker dye. **D)** Quantification of average mitochondrial length, footprint and network in individual chemo-naïve, Doxo-P or Dtx-P MDAMB-231 cells (^****^P < 0.0001, ^***^ <0.001, ^**^<0.01, ^*^<0.05, one-way ANOVA).

Given that the upregulated OXPHOS transcriptional signature of quiescent chemotherapy-persistent TNBC cells included many components of the mitochondrial ETC complexes (**Figure 2A)**, we also examined the respiratory function in this experimental condition. Using the fluorescent dye JC1, which accumulates in the mitochondrial matrix and is used to estimate the outer mitochondrial membrane (OMM) polarization state [35], we found that the OMM potential was higher in quiescent Doxo-P MDAMB-231 cells, compared to Dtx-P and chemo-naïve counterparts (**Figure 2B**). Measurement of oxygen consumption rate (OCR) using a Seahorse analyzer demonstrated that quiescent OXPHOS-high Doxo-P (but not OXPHOS-low Dtx-P) MDAMB-231 cells have significantly increased respiratory function compared to chemo-naïve cells (e.g. **Figure 2C**). This increased OXPHOS function is congruent with the upregulation of OXPHOS genes and the mitochondrial elongation/remodeling in this quiescent cancer cell state. Additional Seahorse-based metabolic analyses in TNBC HC1806 (**Figure S2A)** and SUM159 (**Figure S2B)** cell lines confirmed that Doxo-P (but not Dtx-P) cells had increased respiratory function, supporting that this phenotype is frequent among persistent TNBC cell types. Furthermore, inhibition of OXPHOS using the highly specific OXPHOS inhibitor IACS-010759 [21] rapidly induced apoptosis in quiescent OXPHOS-high Doxo-P (but not in proliferative chemo-naïve or in OXPHOS-low Dtx-P) TNBC cells (e.g. **Figure 2D**) – a finding concordant with previous studies showing that anthracycline-persistent residual tumors acquire dependency on OXPHOS [21].

**Figure 2.**
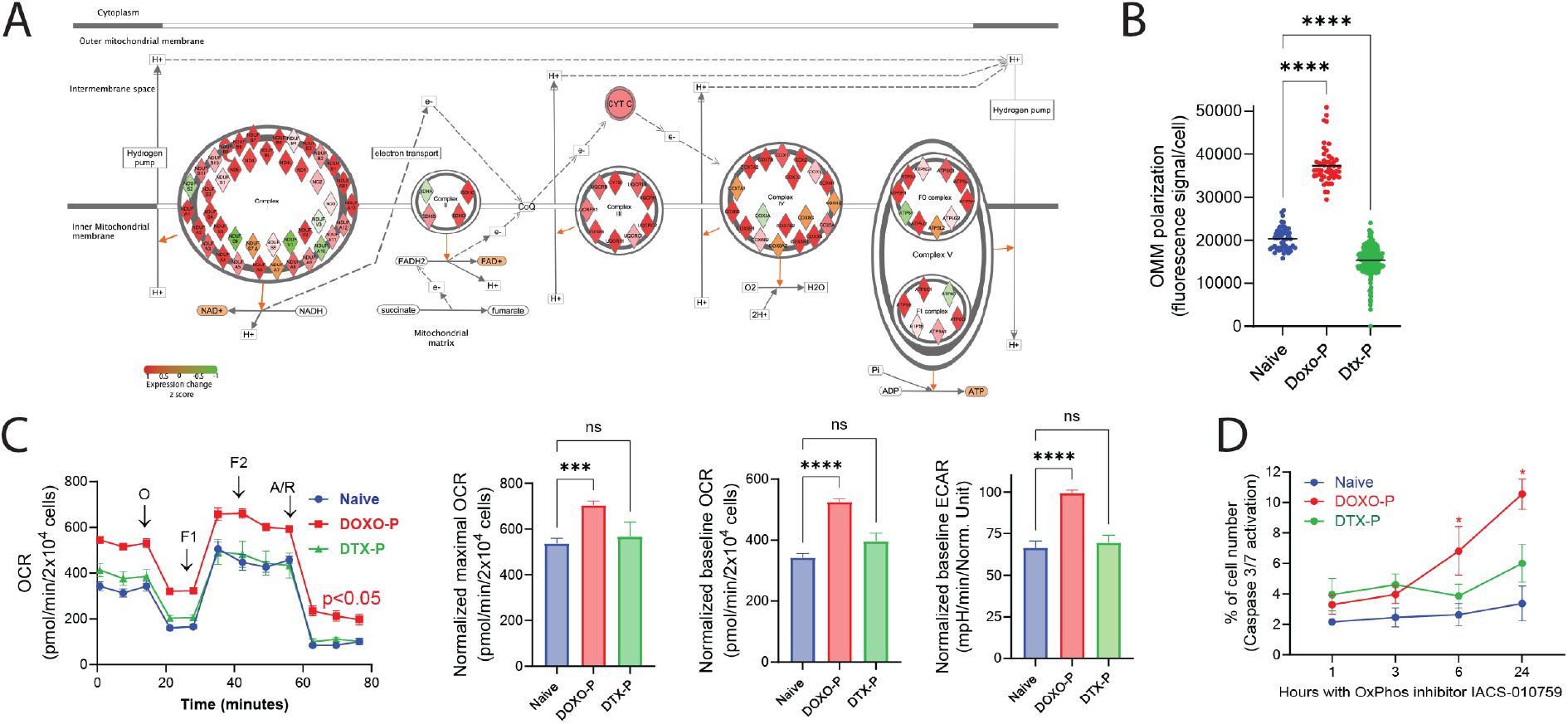
Upregulated mitochondrial respiration in doxorubicin-persistent TNBC cells. **A)** Transcriptional changes (RNA sequencing) in electron transport chain genes in Doxo-P MDAMB-231 cells (compared to chemo-naive). Upregulated genes depicted in red; downregulated in green. **B)** OMM polarization levels in individual cells from chemo-naïve and chemotherapy-persistent MDAMB-231 cells, measured using JC-1 staining assay [35] (one-way ANOVA). **C)** Mitochondrial respiration in chemo-naïve and chemotherapy-persistent MDAMB-231 cells measured using the Seahorse analyzer; values normalized to cell number ((^****^P < 0.0001, ^***^<0.001,^**^<0.01, ^*^ <0.05, one-way ANOVA). **D)** Caspase 3/7 activation in chemo-naïve and chemotherapy-persistent MDAMB-231 cells treated with OXPHOS inhibitor IACS-10759 (1µM); mixed effects analysis.

### Quiescent OXPHOS-high TNBC cells depend on BCL-XL for survival

To determine the therapeutic vulnerabilities of the OXPHOS-high TNBC cells through an unbiased approach, we performed genome-wide CRISPR editing screens in proliferative chemo-naïve and quiescent Doxo-P MDAMB231 cells (schematically depicted in **Figure 3A)**. We quantified the depletion of sgRNAs in chemo-naïve cells (vs. baseline) and in Doxo-P cells (vs. chemo-naïve) to identify the genes that are essential for survival in the quiescent persistent state but not in the proliferative state. *BCL-XL* was the top essential gene for survival in Doxo-P quiescent MDAMB-231 cells (**Figure 3B**). Remarkably, *BCL-XL* was not required for proliferation or survival in chemo-naïve cells (**Figure 3C**). Other notable anti-apoptotic protein genes, such as *BCL2* and *MCL1*, were not essential for survival in Doxo-P cells. The sgRNAs targeting tumor suppressor genes tend to be enriched in chemo-naïve cells (vs baseline) due to the anti-proliferative function of these genes. Consequently, these sgRNAs against tumor suppressor genes were more abundant in proliferating cells than in quiescent cells at the end of the CRISPR editing experiment and therefore scored as depleted in the proliferatively-arrested Doxo-P condition (vs. chemo-naïve). To account for the sgRNAs that score as depleted in Doxo-P (vs. chemo-naïve) due to the quiescent state of Doxo-P cells, we compared the fold changes of all statistically significant (FDR<0.05) depleted sgRNAs in Doxo-P condition (vs. chemo-naïve) with the respective fold changes in chemo-naïve condition (vs. baseline). This comparison confirmed that BCL-XL sgRNA depletion reflected its survival role in Doxo-P cells and not tumor suppressor function in chemo-naive cells (**Figure 3D**). Additional genes that were essential for survival in Doxo-P cells (but did not affect proliferation of chemo-naïve cells) included membrane drug transporter *ABCC1* (also known as multidrug resistance-associated protein 1 [MRP1]), DNA double-strand break repair protein *NHEJ1* and the synthetase of detoxifying molecule selenophosphate *SEPHS2*. The relevance of these additional hits with the mechanism of action of doxorubicin further supports the validity of our CRISPR screen results. The pro-survival function of *BCL-XL* in quiescent Doxo-P cells was confirmed by single-gene knock-out (KO) experiments in TNBC cell lines MDAMB-231, SUM159 and HC1806 (**Figure 3E**). Notably, *BCL-XL* transcript levels were associated with worse outcome in triple-negative (but not in ER-positive) breast cancer patients, supporting the clinical relevance of our findings (**Figure S3**). Interestingly, BCL-XL transcript and protein levels were not upregulated in Doxo-P cancer cells (**Figure 3F, G**). Congruently, qualitative immunohistochemical analysis of paired (pre- and post-treatment) tumor samples from 10 patients treated with anthracycline-based NAC indicated that BCL-XL immunostaining intensity in residual tumors was similar to baseline (representative example shown in **Figure 3H**).

**Figure 3.**
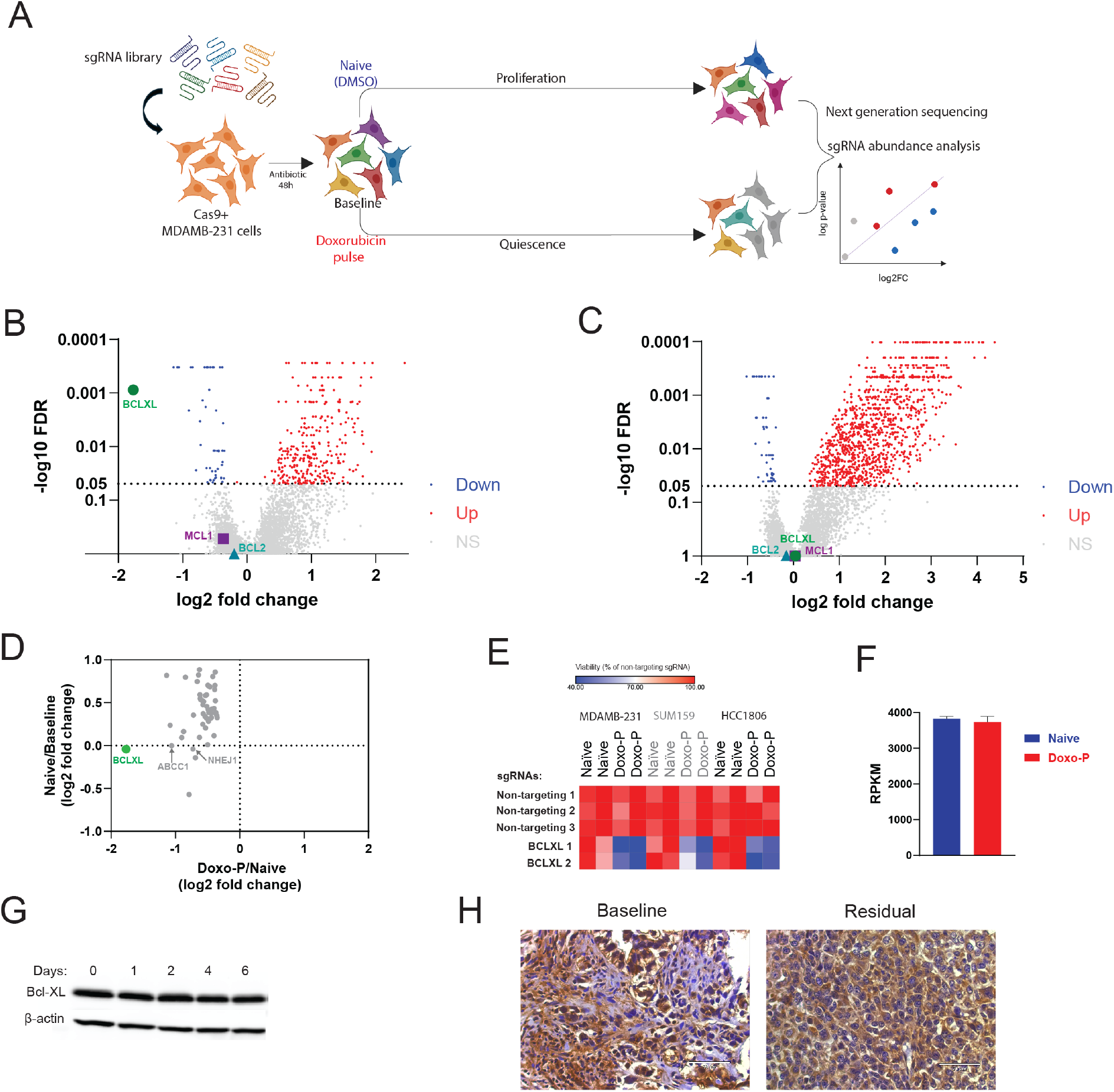
Doxorubicin-persistent cancer cells depend on BCL-XL for survival. **A)** Schematic diagram depicting the genome-wide CRISPR-editing screen in chemo-naïve and Doxo-P MDAMB-231 cells. **B-C)** Changes of sgRNA abundance in doxorubicin-persistent (B) and chemo-naïve (C) MDAMB-231 cells. Analysis performed using the MAGECK pipeline [60]. Genes with significant (FDR<0.05) sgRNA depletion are highlighted in blue or red. **D)** SgRNA fold changes for significantly depleted genes in the Doxo-P vs. chemo-naïve analysis and the respective changes in the chemo-naïve vs. baseline analysis. **E)** Viability of chemo-naïve and Doxo-P MDAMB-231, SUM159 and HCC1806 TNBC cells after *BCL-XL* gene knockout. **F)** *BCL-XL* transcript levels in chemo-naïve and Doxo-P MDAMB-231 cells. **G)** BCL-XL protein levels in MDAMB-231 after treatment with doxorubicin. **H)** BCL-XL protein immunostaining in baseline and post-NAC residual tumor from a TNBC patient (magnification 200x).

### Inhibition of BCL-XL disrupts mitochondrial remodeling and OXPHOS function in Doxo-P TNBC cells

We initially evaluated the apoptotic priming of Doxo-P MDAMB-231 cells using the BH3 profiling method, which is based on longitudinal OMM potential measurement as indication of OMM permeabilization [36-38]. As expected, the OXPHOS-high Doxo-P cells had higher baseline OMM potential compared to their chemo-naïve counterparts. Notably, the Doxo-P cells were sensitive to the apoptotic activator BID (e.g. **Figure 4A** left), indicating that these persistent quiescent cells are apoptotically competent. However, BCL-XL-specific pro-apoptotic sensitizer peptide HRK did not induce apoptosis in Doxo-P cells (**Figure 4A** right), indicating that the protective role of BCL-XL may not necessarily be through its canonical role as negative regulator of pro-apoptotic proteins. Interestingly, previous work in various cell types and biological contexts has demonstrated that BCL-XL can co-localize and interact with mitofusins 1 and/or 2 in the OMM to enable mitochondrial fusion/elongation and apoptosis evasion [39-43]. Concordant with these previous observations, BCL-XL co-immunoprecipitated with MFN2 (but not MFN1) in quiescent Doxo-P (but not in chemo-naïve) MDAMB-231 and SUM-159 TNBC cells (**Figure 4B**). Importantly, 1 hour exposure to the selective BCL-XL inhibitor A-1155463 disrupted mitochondrial elongation and abrogated respiratory function in Doxo-P (but not in chemo-naïve) MDAMB-231 cells, which was followed by caspase activation only after several hours (**Figure 4C, D, E**). These data suggest that BCL-XL may be involved in the mitochondrial remodeling and the upregulation of OXPHOS function in Doxo-P cells, ultimately enabling cell survival.

**Figure 4.**
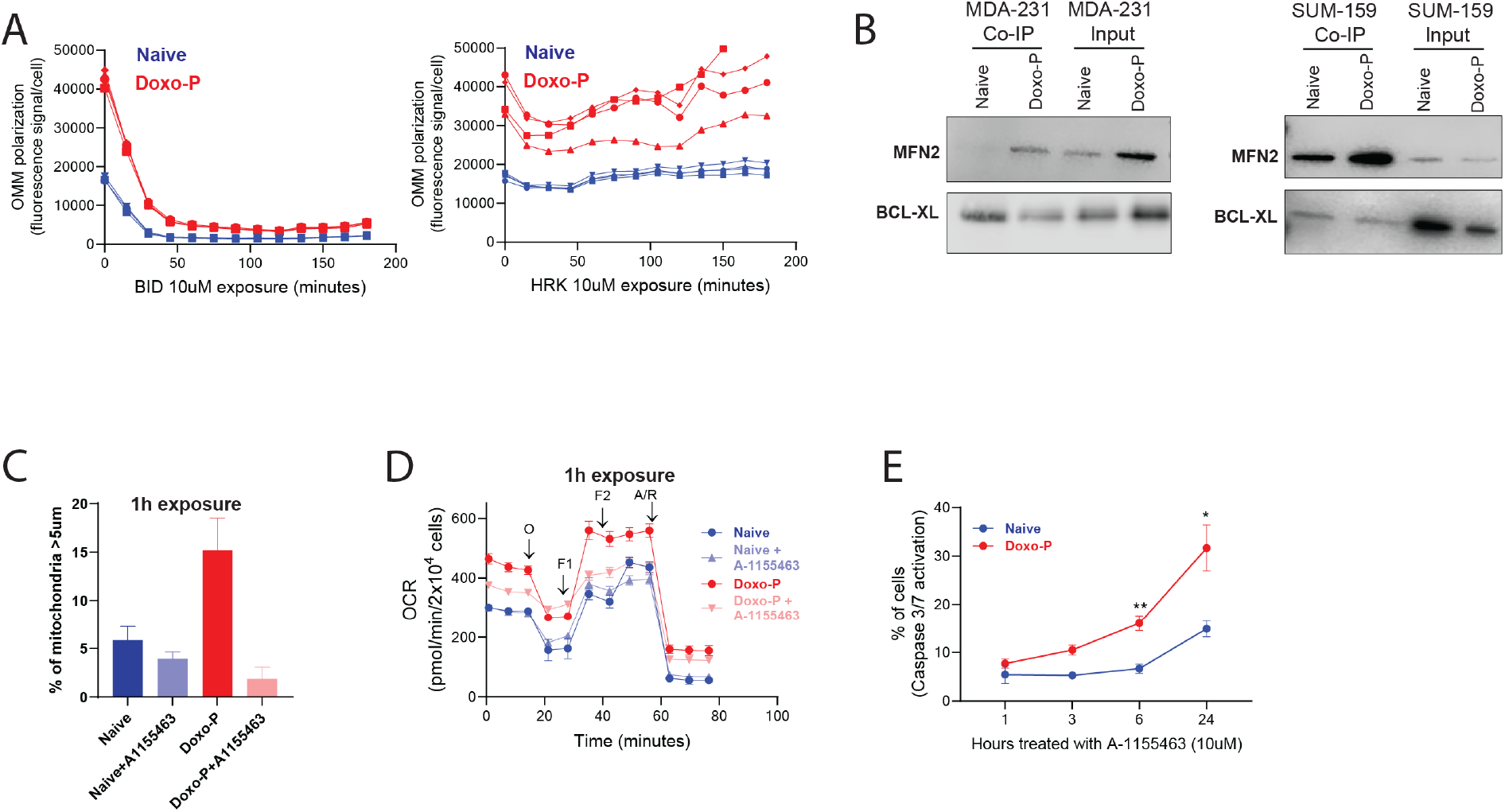
Inhibition of BCL-XL abrogates mitochondrial elongation and respiratory function in doxorubicin-persistent TNBC cells. **A)** The effect of BID and HRK on OMM polarization level in chemo-naïve and Doxo-P MDAMB-231 cells; BH3 profiling assay [38]. **B)** Immunoprecipitation of BCL-XL in chemo-naïve and Doxo-P TNBC cells and analysis by western blot using antibodies against MFN2 and BCL-XL. **C–E)** Effects of BCL-XL inhibitor A-1155463 (10µM, 1h exposure) on mitochondrial length (C), respiratory function measured using Seahorse analyzer (D), and apoptosis (E) in chemo-naïve and Doxo-P MDAMB-231 cells.

### Inhibition of BCL-XL abrogates survival of Doxo-P TNBC cells

Based on our functional genomics results, we hypothesized that therapeutic agents targeting BCL-XL would kill the quiescent anthracycline-persistent TNBC cells. We initially evaluated BCL-XL as therapeutic target in chemo-persistent TNBC cells using the well-validated BCL-XL and BCL2 dual inhibitor navitoclax [44], and the BCL-XL specific inhibitors A1155463 [45] and A1331852 [46] and the cell line models MDAMB-231, HCC1806 and SUM-159. These inhibitors were highly effective against quiescent OXPHOS-high Doxo-P cells, but not against OXPHOS-low Dtx-P or chemo-naïve cells (**Figure 5A, B, C**). Similar results were obtained using the TNBC patient-derived organoid model HCI-002 (**Figure 5D)** that we developed and characterized in our previous studies [1]. Congruent with the CRISPR screen data, the sensitivity of Doxo-P TNBC cells to BCL2 and MCL1 inhibitors was similar to that of chemo-naïve and Dtx-P cells (**Figure S4A)**. Quiescent TNBC cells that persisted after epirubicin or cyclophosphamide treatment were also highly sensitive to BCL-XL inhibitors (**Figure S4B)**, suggesting that the dependency of residual cancer cells on BCL-XL may be generalizable across DNA-damaging chemotherapeutics. Importantly, the cancer cells that exited the Doxo-P quiescent state and resumed proliferation after ∼4 weeks in culture had similar BCL-XL inhibitor sensitivity to chemo-naïve cells (**Figure 5E)**, indicating that the BCL-XL dependency of the OXPHOS-high persistent cancer cell state may be transient.

**Figure 5.**
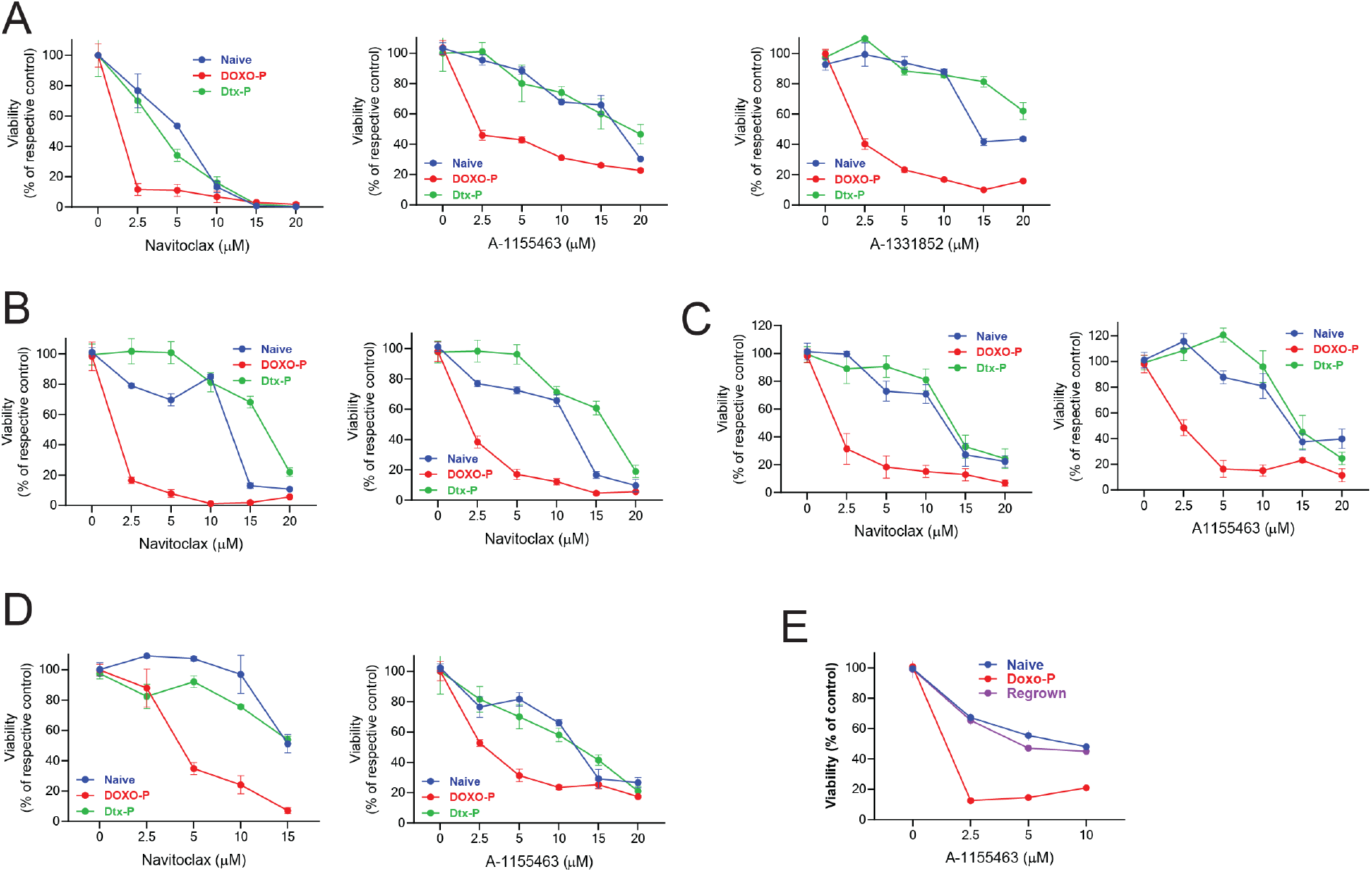
Doxorubicin-persistent TNBC cells are sensitive to inhibitors of BCL-XL. **A-D)** Sensitivity of chemo-naïve, Doxo-P and Dtx-P TNBC cell lines MDAMB-231 (A), HCC1806 (B), SUM159 (C) and TNBC patient-derived organoid line HCI002 (D) to BCL-XL inhibitors navitoclax, A-1155463 and A-1331852. **E)** Sensitivity of chemo-naïve, Doxo-P, and regrown (3–4 weeks post doxorubicin washout) MDAMB-231 cells to Bcl-XL inhibitor A-1155463.

Distinctly from the other TNBC cell models, the quiescent MDAMB-468 TNBC cells that survived after doxorubicin treatment did not have elevated OXPHOS gene expression, mitochondrial elongation or respiratory function (**Figure S5A-C**). Notably, Doxo-P MDAMB-468 did not depend on BCL-XL for survival (**Figure S5D, E**), even though *BCL-XL* gene was expressed in both baseline and Doxo-P conditions (not shown). This observation indicates that the OXPHOS regulation and associated BCL-XL dependency may be heterogeneous among residual Doxo-P TNBC tumors.

Our data suggests that targeting BCL-XL in residual TNBC tumors in combination with anthracyclines can enhance the efficacy of chemotherapy. Existing inhibitors of BCL-XL have potent efficacy against tumor cells but are limited by thrombocytopenia related adverse effects due to the dependence of platelets on BCL-XL for survival [47,48]. To overcome this on-target and dose-limiting toxicity, we recently developed DT2216, a BCL-XL selective proteolysis-targeting chimera (PROTAC) that targets BCL-XL to the Von Hippel-Lindau (VHL) E3 ligase for ubiquitination and subsequent proteasome-mediated degradation [49-51]. Given that VHL is poorly expressed in platelets, DT2216 minimizes platelet loss and is currently being evaluated in phase I/II clinical trials (NCT06620302 & NCT06964009). DT2216 was able to degrade BCL-XL in Doxo-P MDAMB-231 cells, indicating that the proteasome-mediated degradation system is functional in quiescent MYC-suppressed OXPHOS-high cells (**Figure 6A**). As expected, *in vitro* Doxo-P cells were highly sensitive to DT2216, compared to chemo-naïve cells (**Figure 6B**). Combined biweekly treatment of MDAMB-231 xenografts with doxorubicin and DT-2216 (with the first DT2216 dose starting 3 days after the first doxorubicin dose) was superior to either of the two drugs alone (**Figure 6C-H**). Notably, the combination doxorubicin plus DT2216 did not cause significant platelets loss compared to untreated mice (**Figure 6I**), suggesting that this therapeutic combination may avoid the dose-limiting thrombocytopenia observed with other BCL-XL inhibitors.

**Figure 6.**
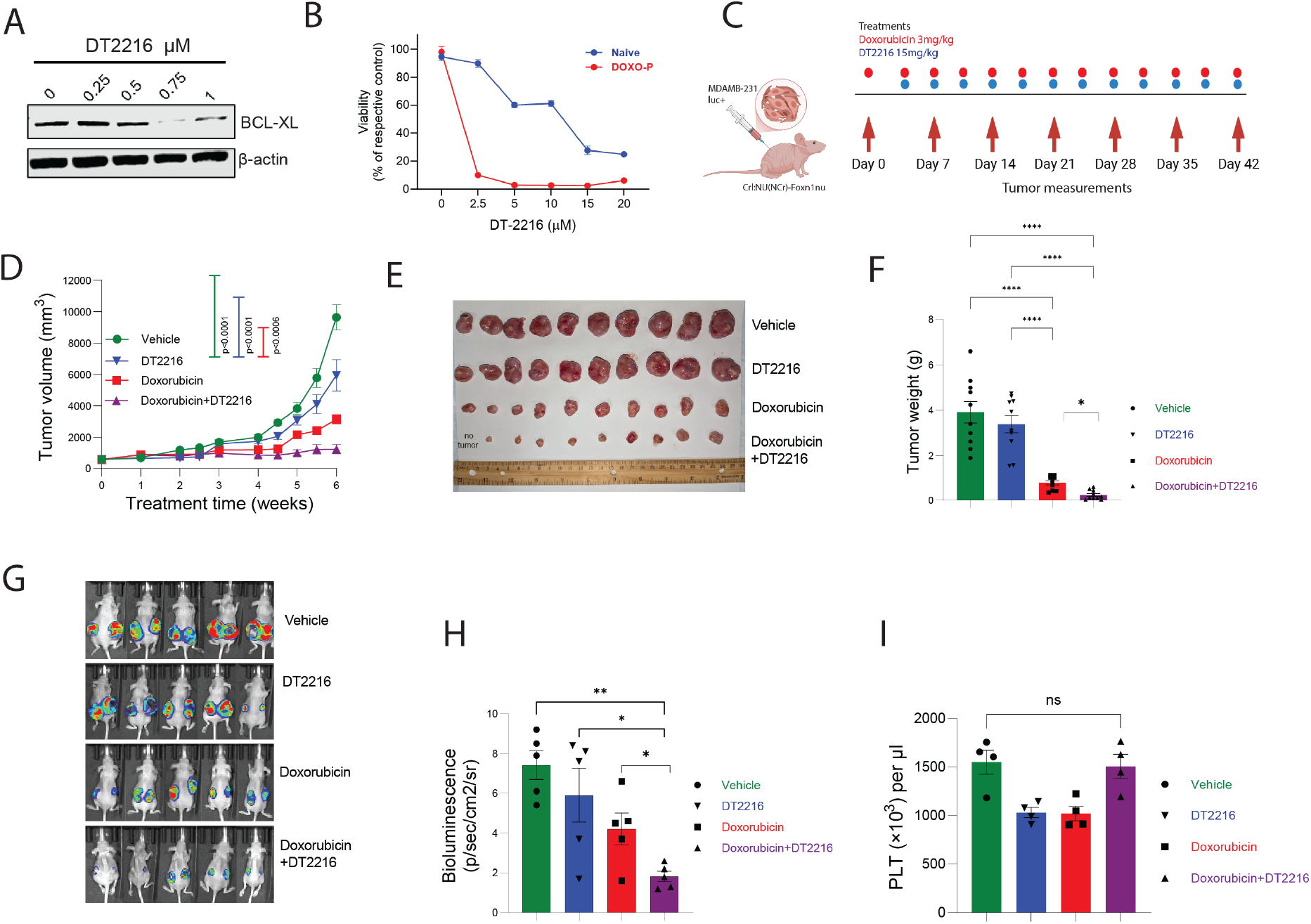
TNBC tumors are sensitive to combination treatment of doxorubicin and DT2216. **A)** DT2216-induced degradation of BCL-XL protein in Doxo-P MDAMB-231 cells. **B)** *In vitro* sensitivity of chemo-naïve and Doxo-P MDAMB-231 cells to DT2216. **C)** Schematic diagram of animal experiment. **D-H)** Response of MDAMB-231 xenografts to combination treatment (twice weekly, IP) of doxorubicin (3mg/kg) and DT2216 (15mg/kg) or each drug alone, estimated by longitudinal measurements of tumor size (D), as well as tumor weight (E-F) and *in vivo* bioluminescence (G-H) after the last drug administration; two-way ANOVA. **I)** Platelet levels in all treatment groups after the last drug administration; two-way ANOVA. ^****^P < 0.0001, ^***^ <0.001, ^**^<0.01, ^*^<0.05.

## Discussion

Chemotherapy is the first-line standard of care treatment for TNBC but often fails to eliminate all malignant cells which can later cause disease relapse. Eradicating the tumor cells that survive the cytotoxic treatment could prevent relapses and prolong remissions. Thus, identifying the mechanisms that enable the survival of persistent cells is critical for the development of more efficient treatment strategies. We and others have shown that tumor cells can survive chemotherapy through a quiescent diapause-like cell state with suppressed global drivers of biosynthesis such as MYC and MTOR [1-3]. Here we find that this diapause-like treatment-persistent cancer cell state is not merely a uniform biosynthetic nadir as consequence of MYC/MTOR suppression, but is actively regulated at transcriptional, metabolic and structural levels in response to distinct chemotherapeutics. Intriguingly, although the expression of mitochondrial OXPHOS genes in proliferating cells requires MYC function [5], OXPHOS was remarkably upregulated in MYC-suppressed anthracycline-persistent TNBC cells. This observation indicates that despite the strongly suppressed MYC transcriptional program, persistent cells can selectively upregulate the OXPHOS gene set through mechanisms that remain to be discovered. Notably, upregulation of OXPHOS has been linked to drug resistance and persistence in several cancers including breast [20], pancreatic [24], lung [4] melanoma [25] and leukemia [23,26]. While the upregulation of OXPHOS genes can explain the increased mitochondrial footprint, we also observed that the mitochondria of Doxo-P cells were elongated and remodeled, confirming previous similar reports [52]. Mitochondrial remodeling could be another mechanism contributing to elevated OXPHOS function in Doxo-P TNBC cells [52]. Distinctly, the TNBC cells that survived treatment with docetaxel did not have upregulated OXPHOS or remodeled mitochondria, indicating that these mitochondrial adaptation mechanisms may not be necessary for persistence to anti-microtubule drugs. These distinct adaptive cell fates upon treatment with anthracyclines and taxanes might have implications regarding the order with which of these two drug classes are administered in chemotherapy regimen.

Accumulating evidence indicates that metabolic rewiring in drug-persistent cancer cells is critical for evading cell death [14,15]. For instance, OXPHOS-high TNBC cells can use glucose or fatty acids as energy sources that contribute to survival [53,54]. Inhibition of OXPHOS, using e.g. ETC complex I inhibitors, can kill drug-resistant cancer cells in many preclinical cancer models [20,21,23-25]. Despite its promise as therapeutic target, direct inhibition of OXPHOS has shown prohibitory systemic toxicity in the clinic [30,31]. We reason that alternative approaches targeting OXPHOS-high persisting TNBC cells could enable the necessary therapeutic index to specifically kill the chemo-persisting cancer cells while sparing normal tissues. Our genomics and pharmacological approaches identified BCL-XL as a key survival dependency in quiescent MYC-low/OXPHOS-high chemotherapy-persistent cancer cell state. Interestingly, only the OXPHOS-high Doxo-P, but not the chemo-naïve or OXPHOS-low Dtx-P, TNBC cells were sensitive to BCL-XL ablation or inhibition, indicating a potential link between the OXPHOS-high state and BCL-XL dependency. The surprising insensitivity of Doxo-P cells to BCL-XL-specific HRK pro-apoptotic peptide turned our attention towards previous reports of a potential role of BCL-XL in mitochondrial remodeling through interactions with mitofusins, which localize in the OMM and are key mediators of mitochondrial fusion [39,40]. Mitochondrial elongation is a complex process that requires coordinated fusion of both inner and outer mitochondrial membranes. For instance, mitochondrial cristae protein OPA1, which mediates inner membrane fusion, is important for the survival of chemotherapy-persistent cancer cells [52]. Our data indicate that BCL-XL interacts with MFN2 and may be involved in OMM fusion, which could explain the rapid mitochondrial fragmentation and collapse of mitochondrial respiration after BCL-XL inhibition in persistent cells. Interestingly, MNF1 and MNF2 did not score as essential for the survival of Doxo-P TNBC cells in the genome-wide CRISPR editing screen, likely reflecting their redundancy for OMM fusion and OXPHOS upregulation in these cells. Our data suggest that BCL-XL-enabled survival of Doxo-P TNBC cells may not necessarily involve its canonical inhibition of pro-apoptotic proteins (e.g. BID/BIM) but could be mediated through a structural role of BCL-XL during mitochondrial aggregation/elongation which contributes to elevated OXPHOS and thus to cancer cell survival. Overall, these findings add on accumulating evidence suggesting that mitochondrial fusion and elongation contribute to drug resistance in breast and other cancers [22,52].

Interestingly, quiescent Dtx-P TNBC cells did not depend on BCL-XL for survival – a finding that seemingly contradicts reports of an anti-apoptotic role of BCL-XL in taxane-treated cells [55]. Crucially, our Dtx-P model reflects a G0/G1 quiescent cell state after drug withdrawal, whereas the anti-apoptotic function of BCL-XL during taxane treatment is exerted during the drug-induced prolonged mitotic arrest and not in non-mitotic cells [55]. Distinctly from anthracycline-persistent cells, the role of BCL-XL during treatment with taxanes suggests that simultaneous (as opposed to sequential) administration of taxanes and a BCL-XL inhibitor may be necessary for the elimination of persistent cells. Additionally, although several Doxo-P TNBC cell lines (MDAMB-231, SUM159 and HCC1806) were OXPHOS-high and dependent on BCL-XL for survival, MDAB-468 Doxo-P cells were OXPHOS-low and were insensitive to BCL-XL ablation or inhibition. Furthermore, Doxo-P cells that exited quiescence lost their sensitivity to BCL-XL inhibitors. These observations highlight the need for predictive tools to assess OXPHOS dependency in residual chemo-persistent tumors, that can be used to identify the patients that would benefit from targeting BCL-XL after chemotherapy, and to correctly time the administration of such treatment.

The management of chemotherapy-induced clinical remission typically involves monitoring for signs of recurrence. Our data support a modification of this clinical practice though administration of sequential “two-hit” strategy to eliminate residual tumors, with the first treatment (e.g. anthracycline) killing a proportion of tumor cells and inducing transition into OXPHOS-high quiescence of the remaining, followed by a second treatment targeting BCL-XL in residual cells and eliminating them. Despite their strong anti-tumor effect, the efficacy of conventional BXL-XL inhibitors was limited by on-target and dose-limiting platelet toxicity, due to the survival dependency of platelets on BCL-XL. On the other hand, thrombocytopenia is also a common adverse effect of anthracyclines, which would render a combination of these two drug classes highly toxic. Here we show that the BCL-XL selective PROTAC DT2216 can be safely combined with doxorubicin to enhance its anti-tumor efficacy. Importantly, this combination did not increase thrombocytopenia rates, indicating that this DT2216 will be better tolerated by patients than other BCL-XL inhibitors. Although this study focuses on TNBC, the derived conclusions are likely to apply to other cancer types treated with anthracyclines and other DNA damaging drugs, therefore potentially impacting a large number of patients.

## Author Contributions

Conceptualization: ED

Methodology: ED, SA, MW, LEG, OA, FMM, EZ, AS, JAG, MK, GZ, SAF, DZ, EG, TW

Investigation: ED, SA, MW, LEG, OA, SAF, TW Visualization: ED, MW, LEG

Funding acquisition: ED Project administration: ED Supervision: ED

Writing – original draft: ED

Writing – review & editing: all authors

## Institutional Review Board Statement

Animal experiments were conducted under animal protocol 00001590, approved by Albert Einstein College of Medicine Institute for Animal Studies (renewed approval on 04/08/2025).

## Data Availability Statement

The datasets used and/or analyzed during the current study are available from the corresponding author on reasonable request.

