## Supplemental Figures for "Quiescent OXPHOS-high triple-negative breast cancer cells that persist after chemotherapy depend on BCL-XL for survival"

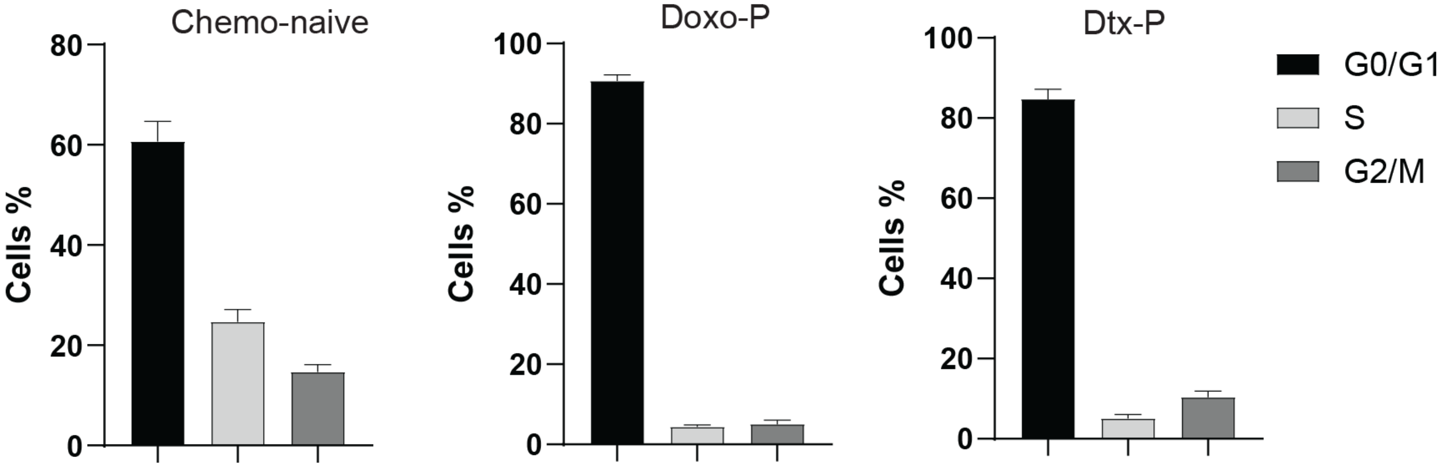


**Supplemental Figure 1**. **Quiescent state of chemotherapy-persistent TNBC cells.** Cell cycle analysis of chemo-naïve and chemotherapy-persistent MDAMB-231 cells 3 days after 2h pulse treatment with indicated drugs.

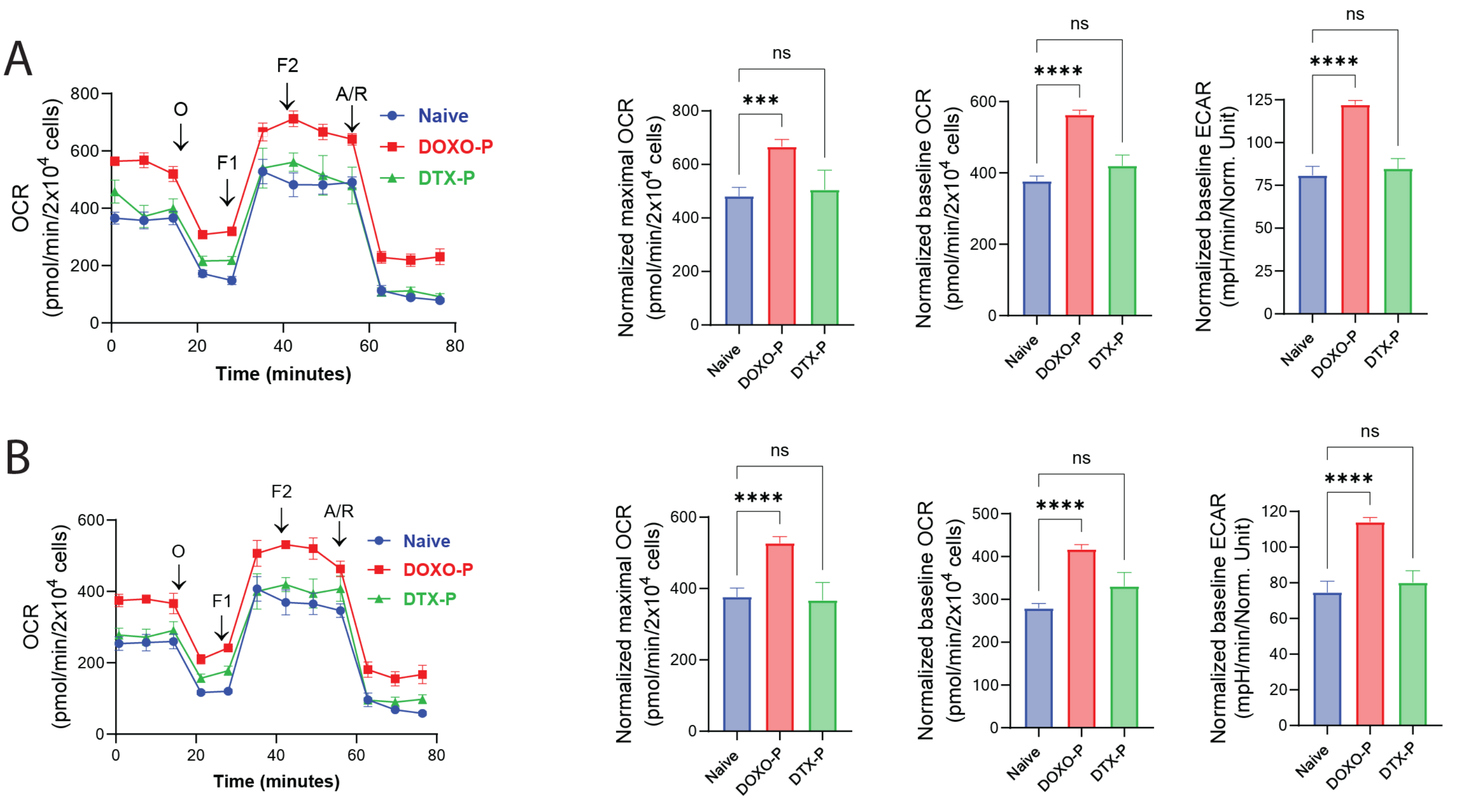


**Supplemental Figure 2**. **Upregulated mitochondrial respiration in doxorubicin-persistent TNBC cells. A-B)** Mitochondrial respiration in chemo-naïve and chemotherapy-persistent HCC1806 (A) and SUM159 (B) TNBC cells measured using the Seahorse analyzer; values normalized to cell number; ****P < 0.0001, ***<0.001, **<0.01, *<0.05, one-way ANOVA.

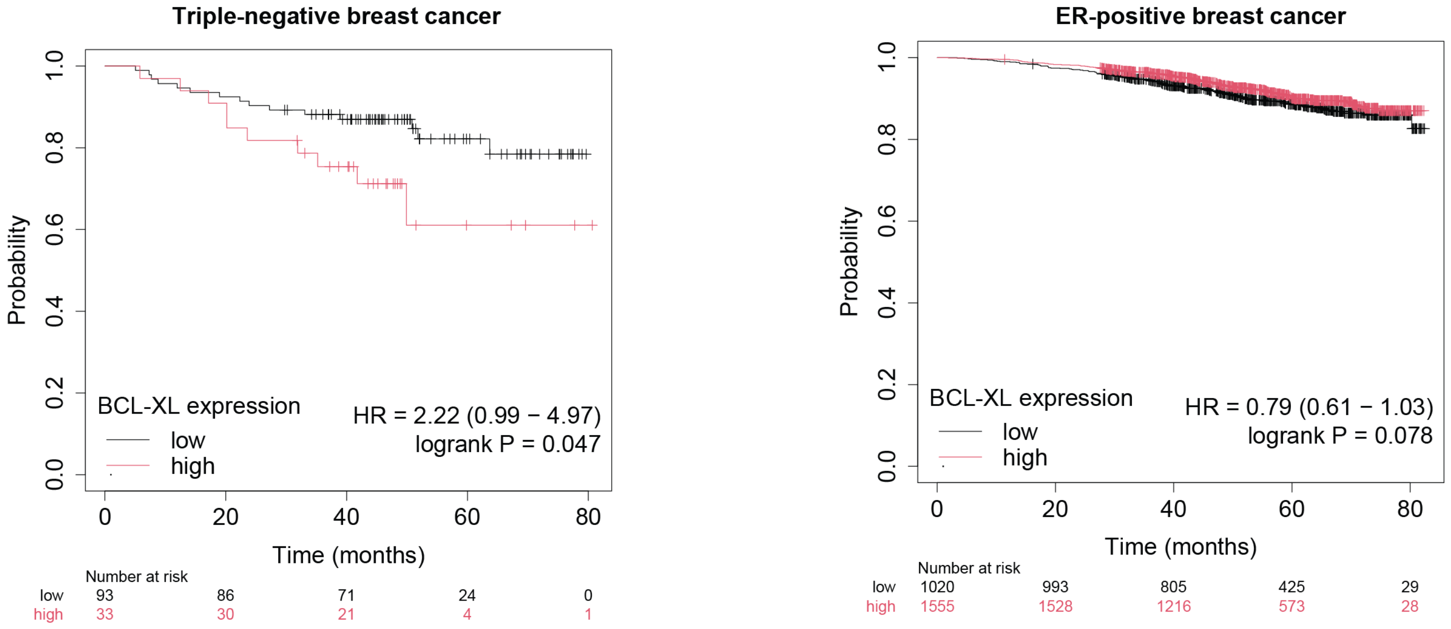


**Supplemental Figure 3**. **Prognostic role of BCL-XL in breast cancer.** Correlation of BCL-XL expression with Overall Survival in triple-negative and ER-positive breast cancer patients. KM-plotter online survival analysis tool [65] accessed on July 31$^{st}$ 2025.

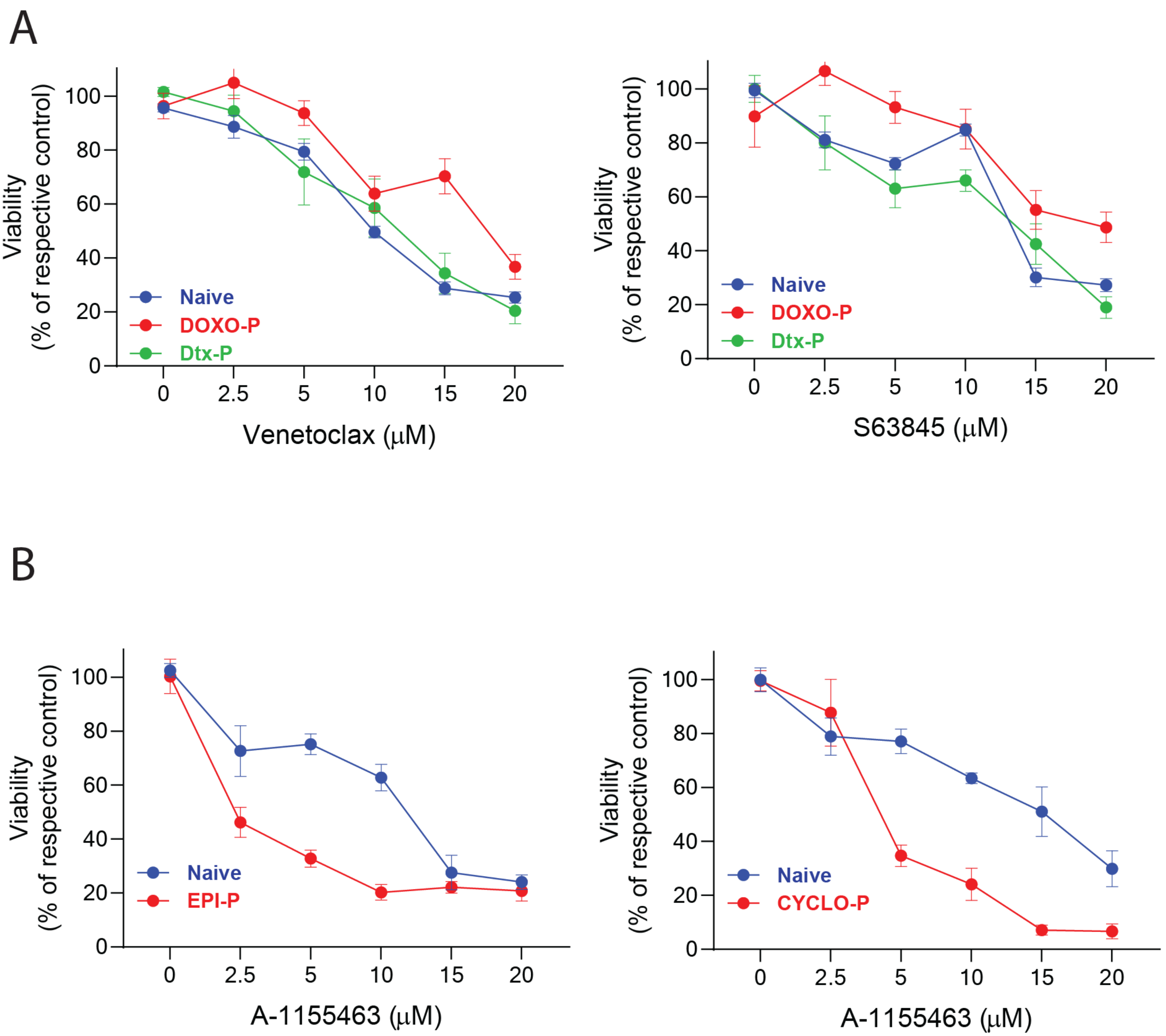


**Supplemental Figure 4**. **Sensitivity of chemotherapy-persistent MDAMB-231 cells to inhibitors of BCL2, MCL1 and BCL-XL. A)** Sensitivity of chemo-naïve, Doxo-P and Dtx-P MDAMB-231 cells to BCL2 inhibitor venetoclax and MCL1 inhibitor S63845. **B)** Sensitivity of chemo-naïve, epirubicin-persistent (Epi-P) and cyclophosphamide-persistent (Cyclo-P) MDAMB-231 cells to BCL-XL inhibitor A1155463.

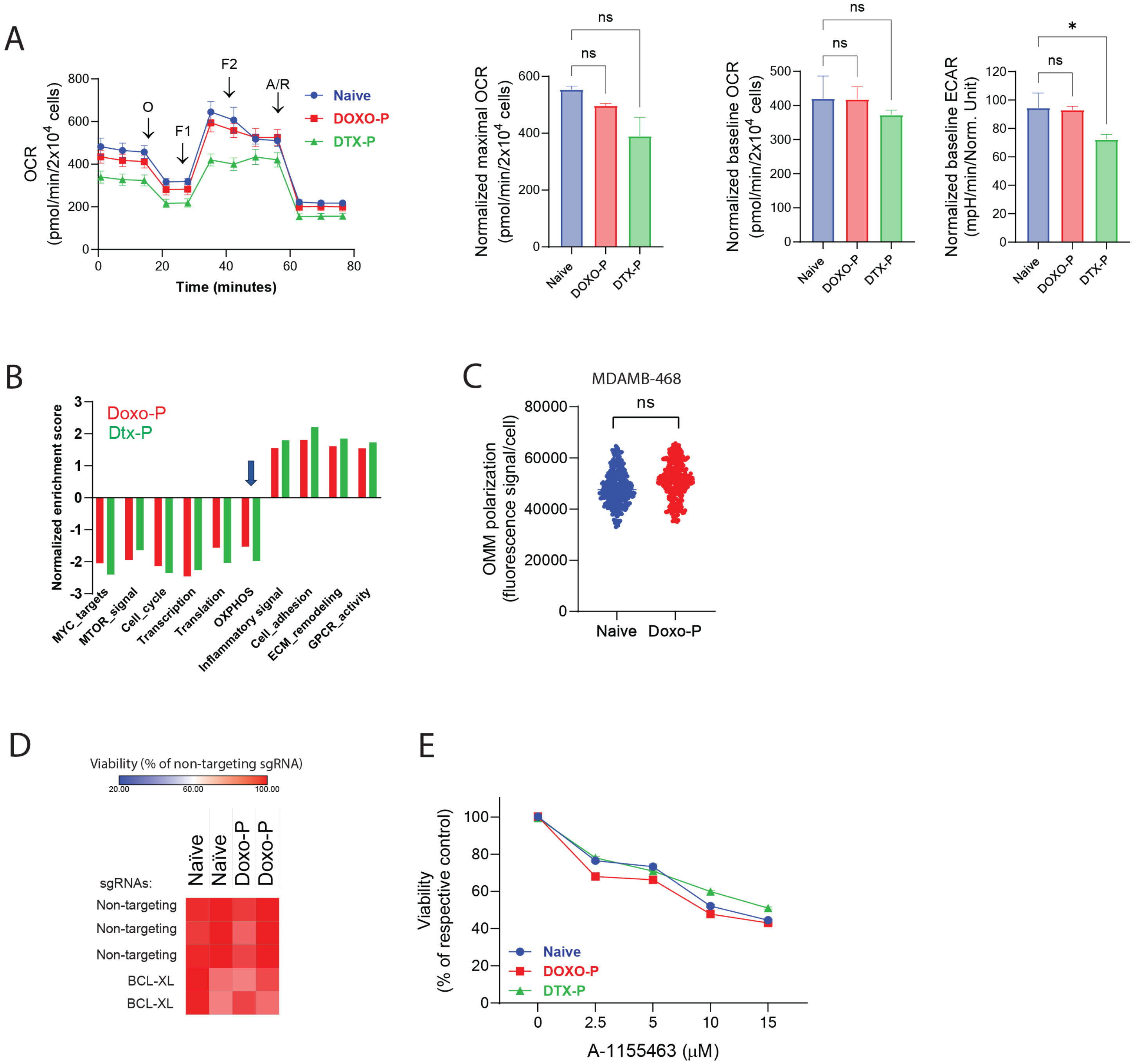


**Supplemental Figure 5**. **Doxorubicin-persistent MDAMB-468 cells have low OXPHOS levels and do not depend on BCL-XL for survival. A)** Mitochondrial respiration in chemo-naïve and chemotherapy-persistent MDAMB-468 cells measured using the Seahorse analyzer; values normalized to cell number. **B)** GSEA analysis of transcriptional changes in MDAMB-468 cells surviving treatment with doxorubicin (Doxo-P) or docetaxel (Dtx-P), compared to chemo-naïve counterparts. **C)** OMM polarization levels in individual cells from chemo-naïve and Doxo-P MDAMB-468 cells, measured using the JC-1 staining assay. **D)** Viability of chemo-naïve and Doxo-P MDAMB-468 after *BCL-XL* gene knockout. **E)** Sensitivity of chemo-naïve, Doxo-P and Dtx-P MDAMB-468 cells to BCL-XL inhibitor A1155463.
